# Enhanced IL-15-mediated NK cell activation and proliferation by an ADAM17 function-blocking antibody involves CD16A, CD137, and accessory cells

**DOI:** 10.1101/2024.05.09.593347

**Authors:** Anders W. Matson, Rob H. Hullsiek, Kate J. Dixon, Sam Wang, Anders J. Lindstedt, Ryan R. Friess, Shee Kwan Phung, Tanya S. Freedman, Martin Felices, Emily N Truckenbrod, Jianming Wu, Jeffrey S Miller, Bruce Walcheck

**Affiliations:** Graduate Program in Comparative and Molecular Biosciences, University of Minnesota, Saint Paul, MN 55108, USA; Graduate Program in Microbiology, Immunology, and Cancer Biology, University of Minnesota, Minneapolis, MN 55455, USA; Department of Veterinary and Biomedical Sciences, University of Minnesota, St. Paul, MN, 55108, USA; Medical Scientist Training Program, University of Minnesota, Minneapolis, MN 55455, USA; Graduate Program in Medicinal Chemistry, University of Minnesota, Minneapolis, MN 55455, USA; Department of Pharmacology, University of Minnesota, Minneapolis, MN 55455, USA; Center for Immunology, University of Minnesota, Minneapolis, MN 55455, USA; Masonic Cancer Center, University of Minnesota, Minneapolis, MN 55455, USA; Department of Medicine, Division of Hematology, Oncology, and Transplantation, University of Minnesota, Minneapolis, MN 55455, USA

## Abstract

**Background:** NK cells are being extensively studied as a cell therapy for cancer. Their effector functions are induced by the recognition of ligands on tumor cells and by various cytokines. IL-15 is broadly used to stimulate endogenous and adoptively transferred NK cells in cancer patients. These stimuli activate the membrane protease ADAM17, which then cleaves assorted receptors on the surface of NK cells as a negative feedback loop to limit their activation and function. We have shown that ADAM17 inhibition can enhance IL-15-mediated NK cell proliferation *in vitro* and *in vivo*. In this study, we investigated the underlying mechanism of this process.

**Methods:** PBMCs or enriched NK cells from human peripheral blood, either unlabeled or labeled with a cell proliferation dye, were cultured for up to 7 days in the presence of rhIL-15 +/- an ADAM17 function-blocking antibody. Different versions of the antibody were generated; Medi-1 (IgG1), Medi-4 (IgG4), Medi-PGLALA, Medi-F(ab′)_2_, and TAB16 (anti-ADAM17 and anti-CD16 bispecific) to modulate CD16A engagement on NK cells. Flow cytometry was used to assess NK cell proliferation and phenotypic markers, immunoblotting to examine CD16A signaling, and IncuCyte-based live cell imaging to measure NK cell anti-tumor activity.

**Results:** The ADAM17 function-blocking mAb Medi-1 markedly increased initial NK cell activation by IL-15. Using different engineered versions of the antibody revealed that the activating Fcγ receptor CD16A, a well-described ADAM17 substrate, was critical for enhancing IL-15 stimulation. Hence, Medi-1 bound to ADAM17 on NK cells can be engaged by CD16A and block its shedding, inducing and prolonging its signaling. This process did not promote evident NK cell fratricide, phagocytosis, or dysfunction. Synergistic activity by Medi-1 and IL-15 enhanced the upregulation of CD137 on CD16A^+^ NK cells and augmented their proliferation in the presence of PBMC accessory cells.

**Conclusions:** Our data reveal for the first time that CD16A and CD137 underpin Medi-1 enhancement of IL-15-driven NK cell activation and proliferation, respectively. The use of Medi-1 represents a novel strategy to enhance IL-15-driven NK cell proliferation, and it may be of therapeutic importance by increasing the anti-tumor activity of NK cells in cancer patients.

**What is already known on this topic:** NK cell therapies are being broadly investigated to treat cancer. NK cell stimulation by IL-15 prolongs their survival in cancer patients. Various stimuli including IL-15 activate ADAM17 in NK cells, a membrane protease that regulates the cell surface density of various receptors as a negative feedback mechanism.

**What this study adds:** Treating NK cells with the ADAM17 function-blocking mAb Medi-1 markedly enhanced their activation and proliferation. Our study reveals that the Fc and Fab regions of Medi-1 function synergistically with IL-15 in NK cell activation. Medi-1 treatment augments the upregulation of CD137 by NK cells, which enhances their proliferation in the presence of PBMC accessory cells.

**How this study might affect research, practice, or policy:** Our study is of translational importance as Medi-1 treatment in combination with IL-15 could potentially augment the proliferation and function of endogenous or adoptively transferred NK cells in cancer patients.

**Graphical abstract:** 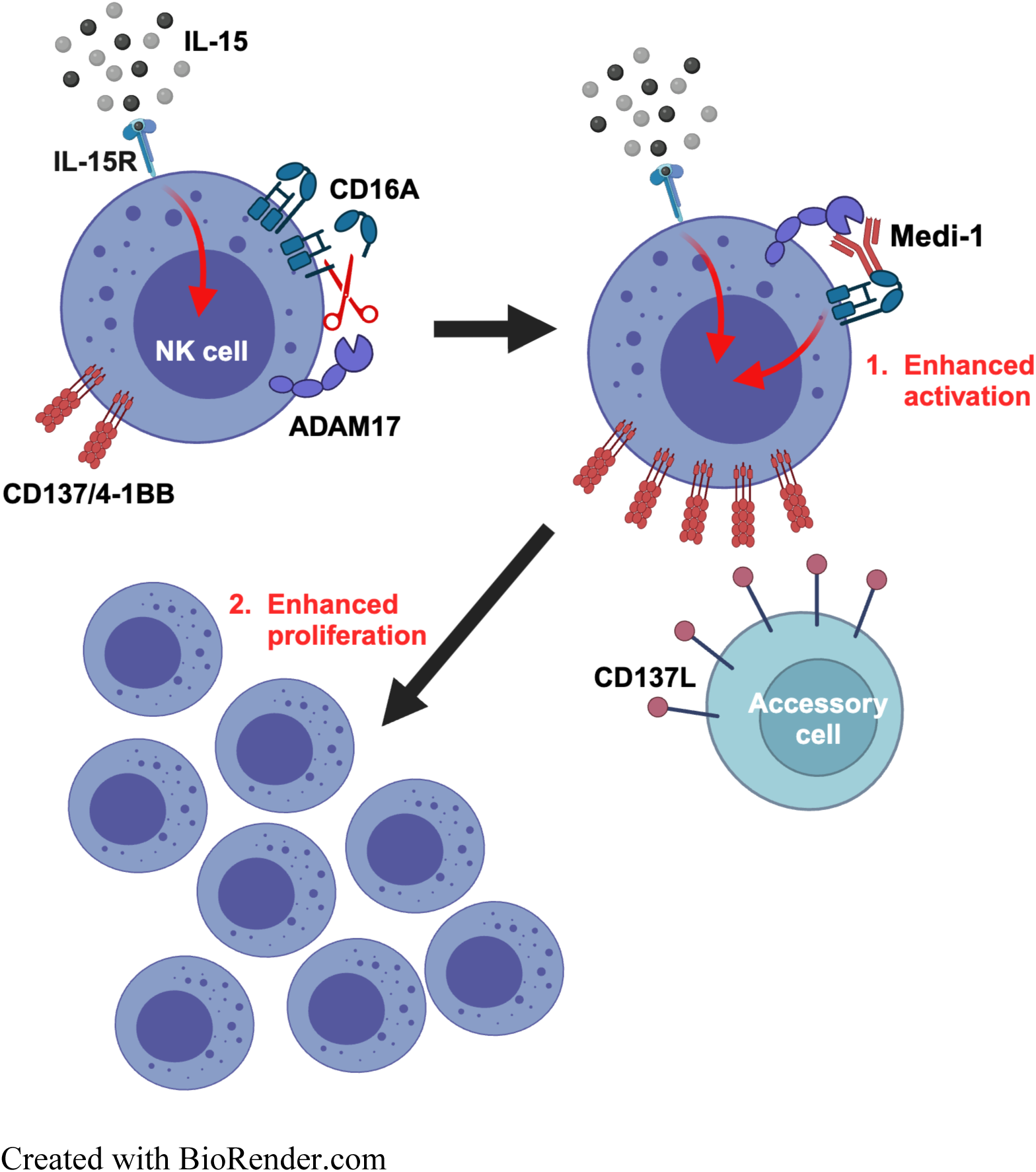

## BACKGROUND

Natural killer (NK) cells are innate lymphocytes that use germline-encoded receptors to interrogate cells of the body for ligands that are either aberrantly upregulated or downregulated to mediate natural cytotoxicity. Human NK cells also target cells by recognizing cell-bound IgG antibodies via their Fc receptor CD16A (FcγRIIIA), inducing antibody-dependent cell-mediated cytotoxicity (ADCC) ^1^. Due to their broad anti-tumor effector functions, there has been a growing emphasis on invoking endogenous NK cells in cancer patients and the adoptive transfer of autologous or allogeneic NK cells ^2–4^. Key challenges in utilizing NK cells for cancer therapies are that they compose a minor population of circulating lymphocytes and their persistence is limited following adoptive transfer ^5 6^. Cytokines such as IL-15 are commonly used to promote human NK cell priming, cytotoxicity, and expansion ex vivo and in patients ^7^.

NK cell activation induces a disintegrin and metalloproteinase-17 (ADAM17) ^8 9^, a membrane-bound protease constitutively expressed on the surface of NK cells that has many substrates ^10 11^. Their cleavage typically occurs in a *cis* manner at an extracellular location proximal to the cell membrane, referred to as ectodomain shedding ^10^. CD16A on natural killer cells is a well-described ADAM17 substrate, which is rapidly cleaved following NK cell activation, as a negative feedback mechanism, and upon its engagement of cell-bound antibodies, as a detachment process ^12^.

We have reported that blocking ADAM17 function with a monoclonal antibody (mAb) markedly enhanced IL-15-driven NK cell expansion ex vivo and in vivo ^9^. In this study, we sought to elucidate the underlying mechanism of this process. We show that Medi-1, a human IgG1 mAb that selectively blocks ADAM17 function, augmented IL-15-driven NK cell proliferation and this involved its Fc region. Our data reveals that Medi-1 is engaged by the activating receptor CD16A and blocks its shedding, which synergizes with IL-15 signaling to enhance CD137 upregulation and NK cell proliferation when in the presence of PBMC accessory cells.

## METHODS

### Antibodies

The anti-human ADAM17 mAb MEDI3622 has been previously described ^13^. The following Fc variants were generated from its nucleotide sequence. Human isotypes IgG1 and IgG4 were generated by Syd Labs (Natick, MA) and are referred to as Medi-1 and Medi-4, respectively. The mutations Pro329Gly, Leu234Ala, and Leu235Ala (PGLALA) were engineered into the Fc region of Medi-1 (Syd Labs) to impair FcγR binding and is referred to as Medi-PGLALA. The Fc region of Medi-1 was proteolytically removed (SouthernBiotech, Birmingham, AL) and is referred to Medi-F(ab′)_2_. The Trispecific Killer Engager (TriKE) containing a human IL-15 moiety and humanized camelid single-domain antibodies against CD16A and B7H3 (cam16-IL15-camB7H3), as well as a Targeted ADAM17 Blocker-16 (TAB16), containing a single-chain fragment variable (scFv) from Medi-1 and a humanized camelid single-domain antibody against CD16, were generated similar to as previously described ^14^. All commercially available antibodies are listed in **Table 1**. Any antibodies used in functional assays that contained a preservative, such as sodium azide (NaN_3_), were subjected to a Zeba Spin Desalting Column 40K (Thermo Fisher Scientific, Waltham, MA) according to the manufacturer’s instructions, and resuspended in PBS buffer without Ca^+2^ and Mg^+2^ (Gibco, Waltham, MA, USA).

**Table 1.** Commercial antibodies.

| ANTIBODY | FLUORESCENCE CONJUGATE | ANIMAL ORIGIN | SPECIES RECOGNIZED | ANTIGEN/LIGAND RECOGNIZED | SOURCE, CATALOG # |
| --- | --- | --- | --- | --- | --- |
| CD56 | Pe/Cy7 | Mouse | Human | CD56 | Biolegend, 318318 |
| CD107a (LAMP-1) | APC | Mouse | Human | CD107a | Biolegend, 328620 |
| CD69 | PerCP/Cy5.5 | Mouse | Human | CD69 | BioLegend, 310927 |
| CD16 | BV510 | Mouse | Human | CD16 | BioLegend, 302048 |
| NKG2D | BV605 | Mouse | Human | NKG2D | BioLegend, 320832 |
| NKp46 | BV650 | Mouse | Human | NKp46 | BioLegend, 331927 |
| KIR2DL1 | FITC | Mouse | Human | KIR2DL | BioLegend, 339504 |
| NKG2A | PE | Mouse | Human | NKG2A | BioLegend, 375104 |
| CD3 | BV785 | Mouse | Human | CD3 | BioLegend, 300472 |
| CD3 | FITC | Mouse | Human | CD3 | BioLegend, 300440 |
| KIR2DL2/L3 | FITC | Mouse | Human | KIR2DL2/L3 | BioLegend, 312604 |
| KIR3DL1 | PE | Mouse | Human | KIR3DL1 | BioLegend, 312708 |
| NKp44 | APC | Mouse | Human | NKp44 | BioLegend, 325110 |
| LAG-3 | BV510 | Mouse | Human | LAG-3 | BioLegend, 369318 |
| TIGIT | BV605 | Mouse | Human | TIGIT | BioLegend, 372712 |
| TIM-3 | BV650 | Mouse | Human | TIM-3 | BioLegend, 345028 |
| CD25 | FITC | Mouse | Human | CD25 | BioLegend, 302604 |
| BBK-2 | NA | Human | Human | CD137 | Invitrogen, MA5-13739 |
| CD137 | APC | Mouse | Human | CD137 | BioLegend, 309810 |
| Human 4-1BBL/CD137L/TNFSF9 ECD, Fc-fusion (sCD137L/Fc) | NA | Human | Human | CD137 | G&P Bioscience, FCL2590B |
| IFN-γ | PE | Mouse | Human | IFN-γ | BioLegend, 502508 |
| CD4 | FITC | Mouse | Human | CD4 | BioLegend, 317408 |
| CD8 | PE | Mouse | Human | CD8 | BioLegend, 344706 |
| Ghost Dye Red 710 | Red 710 |  |  |  | Tonbo Biosciences, 13-0871 |
| D1(A12) | NA | Human | Human | ADAM17 | Merck Millipore Sigma, MABT884 |
| Human IgG1 | NA | Human | NA | NA | Merck Millipore Sigma, AG502-M |
| Urelumab | NA | Human | Human | CD137 | Invitrogen, MA5-141939 |
| Daratumumab | NA | Human | Human | CD38 | Janssen Biotech, Inc |
| pZAP-70/pSYK | NA | Rabbit | Human | Phospho-Zap-70 (Tyr319)/Syk (Tyr352) (65E4) | Cell Signaling Technology, 2717S |
| Total SYK | NA | Mouse | Human | Syk (4D10) | Cell Signaling Technology, 80460 |
| β-actin | NA | Mouse | Human | β-Actin (8H10D10) | Cell Signaling Technology, 3700 |
| anti-Rabbit IgG | IRDye® 800CW | Donkey | Rabbit | IgG | LICORbio, 926-32213 |
| anti-Mouse IgG | IRDye® 680RD | Donkey | Mouse | IgG | LICORbio, 926-68072 |

### Flow Cytometric Analyses

NK cell phenotypic analyses were performed as previously described ^9^. Briefly, fluorescence minus one (FMO) and appropriate isotype-matched antibodies were used for controls. An FSC-A/SSC-A plot was used to set an electronic gate on leukocyte populations, and an FSC-A/FSC-H plot was used to set an electronic gate on single cells. The cell viability dye Ghost Dye Red 710 (Tonbo Bioscience, San Diego, CA) was used to distinguish live vs. dead cells per the manufacturer’s instructions. Cell samples were acquired on a BD FACSCelesta Cell Analyzer (BD Biosciences, San Jose, CA) and analyzed using FlowJo v10.9.0 Software (BD Bioscience).

### Cell isolation and culture

Peripheral blood mononuclear cells (PBMCs) were obtained from whole blood, collected from healthy consenting adults at the University of Minnesota (IRB protocol # 9708M00134) in sodium heparin tubes (BD Bioscience), and from plateletpheresis products from Innovative Blood Resources (St. Paul, MN). PBMCs were isolated using Lymphocyte Separation Medium (Promocell, Heidelberg, Germany). NK cells were enriched using a negative selection human NK Cell Isolation Kit (Miltenyi Biotec, Bergisch, Germany) or (StemCell Technologies, Vancouver, BC, Canada). Isolated NK cells were ≥90% pure, as determined by CD56^+^ CD3^−^ staining and flow cytometry. PBMCs or enriched NK cells, either unlabeled or labeled with CellTrace Violet (CTV) Cell Proliferation Dye (ThermoFisher Scientific) as per the manufacturer’s instructions, were cultured for up to 7 days at 37°C in 5% CO_2_ in RPMI 1640 media (Gibco) supplemented with 10% Heat Inactivated FBS (Gibco) and 1x Anti-Anti (Gibco), in the presence or absence of rhIL-15 or rhIL-2 (R&D Systems, Minneapolis, MN) and the indicated antibodies. NK cell proliferation was analyzed via flow cytometry and a division index was calculated using FlowJo. SKOV-3 cells (HTB-77), an ovarian adenocarcinoma cell line, K562 cells (CCL-243), a chronic myelogenous leukemia cell line, and NK-92 cells (CRL-2407), a malignant non-Hodgkin’s lymphoma NK cell line, were purchased from ATCC (Manassas, VA, USA). SKOV-3 cells stably expressing NucLightGreen (SKOV-3/NLG) were generated as previously described ^15^. SKOV-3 cells were maintained in McCoy’s 5A media (Gibco) supplemented with 10% HI FBS and 1X Anti-Anti. K562 cells were maintained in 10% RPMI 1640 media, as described above. NK-92 cells were maintained in MEM Alpha media (Gibco) with 12.5% HI FBS (Gibco), 12.5% Horse Serum (Gibco), 1x Anti-Anti, and 0.1mM 2-Mercaptoethanol (Gibco). All cell lines were routinely tested for mycoplasma with the MycoAlert Mycoplasma test kit (Lonza, Basel, Switzerland).

### Cellular assays

PBMCs were cultured with SKOV-3 or K562 cells at an effector:target (E:T) ratio of 1:1 at 37°C and 5% CO_2_ for 5 hours. CD107a and IFN-γ staining were determined as previously described ^8^. To measure cell cytotoxicity in real-time, SKOV-3/NLG cells were plated at a density of 4x10^3^ cells/well in a 96-well flat bottom, tissue culture-treated plate (Falcon, cat# 353072, Corning, NY) 24 hours prior to starting the assay. Enriched NK cells were added at the indicated E:T ratios in RPMI 1640 media at 37°C in 5% CO_2_. Fluorescent images of live cells were obtained hourly for 7 days using an IncuCyte SX3 live cell imaging and analysis system (Sartorius Göttingen, Germany), as previously described ^15^.

NK-92 cells expressing human CD16A were stimulated with phorbol 12-myristate 13-acetate (PMA), as described previously ^16^. Briefly, NK-92/CD16A cells (3x10^5^/ml) were treated with or without 10ng/ml PMA (Sigma Aldrich, St. Louis, MO) and 1μg/ml ionomycin (AdipoGen Life Sciences, San Diego, CA) for 2 hours at 37°C in the presence or absence of Medi-1, Medi-4, Medi-PGLALA, Medi-F(ab′)_2_, D1(A12), or an isotype-matched negative control antibody at the indicated concentrations. The cells were then washed twice with flow buffer, treated with cell viability dye, and stained for CD16A. All samples were fixed with 2% paraformaldehyde in PBS without Ca^+2^ and Mg^+2^ and subjected to flow cytometric analyses.

### Immunoblotting

Enriched NK cells were untreated or treated with Medi-1 or an isotype negative control antibody (5μg/ml) for the indicated time points at 37°C in serum-free RPMI 1640 (Gibco). After stimulation cells were washed, detergent lysed, and prepared for immunoblotting as previously described ^17^. Briefly, ≈2.5x10^5^ cells were run in each lane of a 7% NuPAGE tris-acetate gel (Invitrogen, Carlsbad, CA) and transferred to Immobilon-FL polyvinylidene difluoride membrane (EMD Millipore, Burlington, MA). REVERT Total Protein Stain (LI-COR Biosciences, Lincoln, NE) was used as a standard for total protein normalization. Membranes were treated with Intercept (TBS) Blocking Buffer (LI-COR Biosciences), incubated with the indicated primary antibodies, and then incubated with the appropriate near-infrared secondary antibodies. Blots were visualized using an Odyssey CLx near-infrared imager (LI-COR Biosciences), and signals were background-subtracted using ImageStudio Software (LI-COR Biosciences) and corrected for whole-lane protein content (total protein stain).

### Statistical Analysis

Data are presented as the mean ± SD. Compiled *in vitro* data are from at least three independent experiments using separate donors. Statistical analysis was conducted, and significance was determined using GraphPad Prism version 10.0.3 for IOS, GraphPad Software (La Jolla, CA). Comparison between two treatments was computed using the paired two-tailed Student’s t-test. Comparison among three or more treatments was made using one-way or two-way ANOVAs, as appropriate, followed by the indicated post hoc test. All data presented as geometric mean fluorescent intensity (MFI) were log-transformed before statistical analysis. Area under the curve (AUC) was calculated to compare the killing kinetics of the live imaging *in vitro* killing assays. The data generated in this study are available upon request from the corresponding author.

## RESULTS

### Medi-1 treatment selectively amplifies NK cell proliferation in the presence of IL-15

It is well established that IL-15 promotes the proliferation of NK cells and T cells, especially in the presence of monocytes which transpresent IL-15 ^18^. To determine the effects of the ADAM17 function-blocking mAb Medi-1 on IL-15-driven proliferation of these populations, PBMCs labeled with CTV were cultured with IL-15 at 10 ng/mL, a standard concentration used for *ex vivo* lymphocyte expansion ^9 19^, +/- Medi-1 (5 μg/ml) for up to 7 days. As shown in **Figure 1A**, IL-15 stimulation along with Medi-1 was observed to markedly enhanced NK cell proliferation, which was dose-dependent (**Supplemental Fig. 1A**), whereas Medi-1 treatment alone had a negligible effect on NK cell proliferation (**Fig. 1A**). An isotype-matched control for Medi-1 did not augment IL-15-driven NK cell proliferation (**Supplemental Fig. 1B**). Medi-1 treatment also increased NK cell proliferation at a low concentration of IL-15 (1 ng/mL) (**Supplemental Fig. 1C**). Enhanced IL-15-driven NK cell proliferation by Medi-1 caused a corresponding increase in the proportion of these cells in the PBMC population after 7 days of culture (**Fig. 1B**), resulting in a 4.5-fold expansion, with no effect on cell viability, compared to a 1.5-fold expansion exhibited by cells stimulated with IL-15 alone (**Fig. 1C**). In contrast, T cells within the IL-15-stimulated PBMC population were only marginally affected by Medi-1 treatment, with NKT cells showing minor augmentation of proliferation while CD4 and CD8 T cells showed no benefit (**Fig. 1D**). Hence, the Medi-1 effect was primarily NK cell-specific and it increased their sensitivity to IL-15 stimulation.

**Figure 1.**
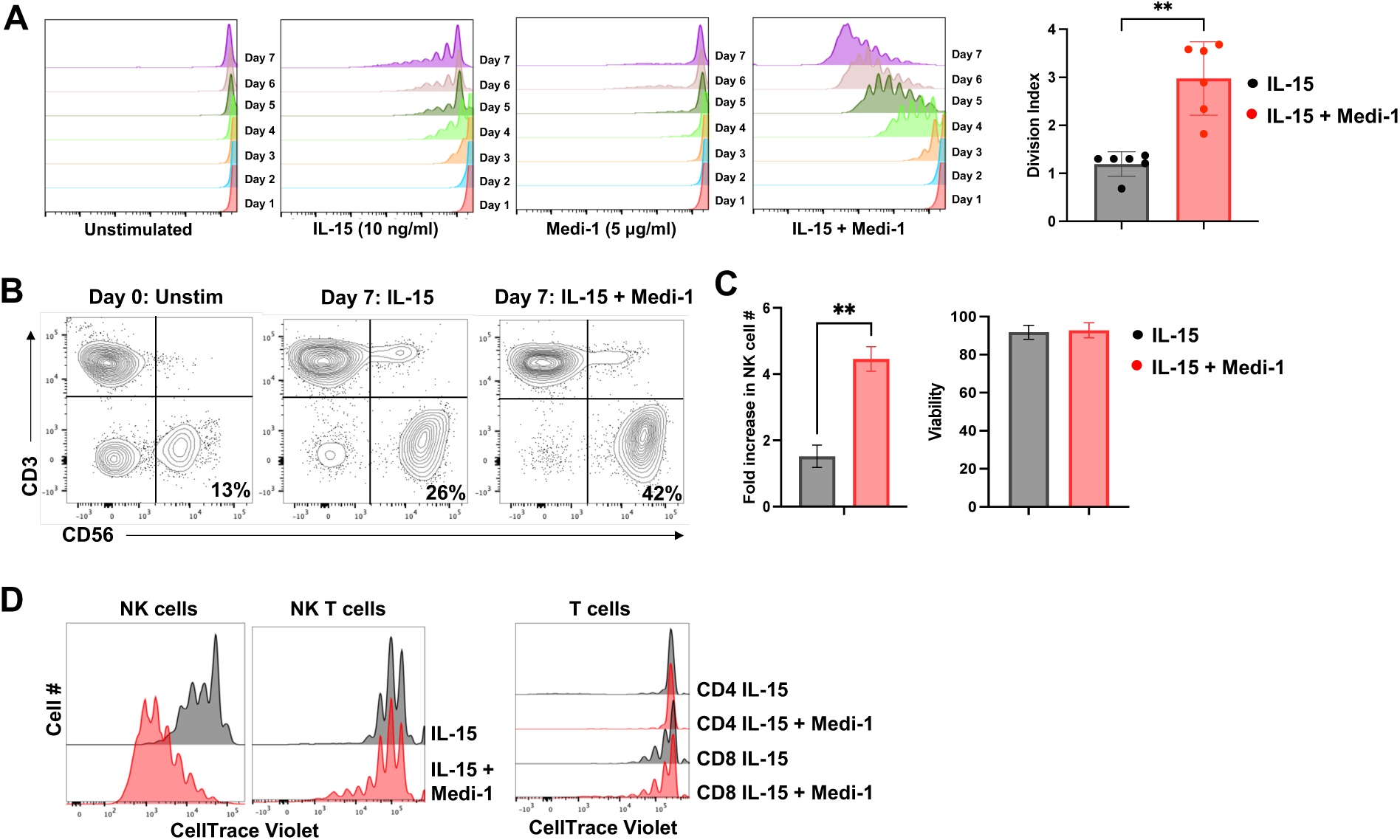
Medi-1 treatment selectively increases NK cell proliferation by IL-15. Freshly isolated PBMCs were labeled with CellTrace Violet (CTV) to track cell division and cultured for 7 days +/- IL-15 (10ng/ml) +/- Medi-1 (5µg/ml). **A.** CD56^+^ CD3^−^ NK cells were analyzed for CTV dilution by flow cytometry. Representative data are shown (left panels). Division index (right panel), mean +/- SD, n=6 donors. **B.** Percentage of NK cells for the indicated conditions and time points. Data are representative of 6 independent experiments using leukocytes from separate donors. **C.** Fold increase in NK cells and viability after 7 days of expansion for the indicated conditions. Mean +/- SD, n=4 donors. **D.** The indicated lymphocyte populations were analyzed for CTV dilution by flow cytometry. NK cells (CD56^+^ CD3^−^), NK T cells (CD56^+^ CD3^+^), CD8 T cells (CD56^−^ CD3^+^ CD8^+^) and CD4 T cells (CD56^−^ CD3^+^ CD4^+^). Data comparing NK cells to T cells are representative of at least 3 independent experiments using leukocytes from separate donors. **p < 0.01; ns=not significant. Statistical significance was determined by paired two-tailed Student’s t-tests.

D1(A12) is another human IgG1 mAb that blocks ADAM17 function ^20^ (**Supplemental Fig. 2**). Interestingly, D1(A12) treatment of PBMCs marginally but did not significantly enhance NK cell proliferation by IL-15 stimulation (**Supplemental Fig. 3**). This could be due to distinct aspects of the Fab and/or Fc regions of D1(A12) or its targeted epitopes. The variable heavy and light chains of this engineered mAb were designed to recognize distinct epitopes in different regions of ADAM17 ^20^. Moreover, the strength of CD16A binding to IgG1 is affected by glycosylation and amino acid polymorphisms in its Fc region ^21^, which could vary between D1(A12) and Medi-1.

IL-15 is used therapeutically in various forms, including its incorporation into multi-engager complexes, such as a Trispecific Killer Engager (TriKE) ^22^. To assess whether Medi-1 treatment enhanced NK cell proliferation by IL-15 in other contexts, we used it in conjunction with a TriKE consisting of a humanized camelid nanobody against CD16, IL-15, and a camelid nanobody against the tumor-associated antigen B7-H3, referred to as cam16-IL15-camB7H3. We observed that Medi-1 treatment enhanced NK cell proliferation mediated by the TriKE (**Supplemental Fig. 1D**). Moreover, the effects of Medi-1 were not restricted to IL-15, as the presence of Medi-1 enhanced IL-2-driven NK cell proliferation as well (**Supplemental Fig. 1E**).

### NK cells undergoing IL-15-driven proliferation in the presence or absence of Medi-1 express similar functional markers

Overstimulation of NK cells with IL-15 or other stimuli can cause changes in their phenotypic profile, including the downregulation of activating receptors and the upregulation of inhibitory receptors ^23^. Stimulation of NK cells with a single 10 ng/ml dose of IL-15 for 7 days does not induce their exhaustion ^19^. To assess whether this is altered by Medi-1 treatment, PBMCs were stimulated in this manner for 7 days +/- Medi-1 and then NK cells were compared for their staining levels of well-described activating and inhibitory receptors by flow cytometry. As shown in **Figure 2A**, we observed no statistical differences in the percentage of NK cells expressing NKp44, NKp46, NKG2D, KIR2DL1, KIR2DL2/L3, KIR3DL1, or NKG2A when stimulated with IL-15 +/- Medi-1. The percentage of NK cells expressing CD16A, an ADAM17 substrate ^10^, was slightly increased in the presence of Medi-1 (**Fig. 2A**). The geometric MFI of positively staining NK cells for NKG2D, KIR2DL1, KIR2DL2/L3, and NKG2A was also marginally increased for IL-15-stimulated NK cells in the presence of Medi-1 (**Fig. 2A**). The inhibitory receptors LAG-3, TIGIT, and TIM-3 have been reported to be upregulated in expression and activity in exhausted T cells ^23 24^. These markers can be upregulated on activated NK cells, but whether their expression signifies exhaustion is less clear ^23^. LAG-3, TIGIT, and TIM-3 staining levels on resting NK cells varied, therefore we included FMO controls in the flow cytometric plots to better distinguish changes in their staining during NK cell activation. When NK cells were cultured in the presence of IL-15 and Medi-1, the percentage of LAG-3^+^ cells was significantly increased, as was the geometric MFI of cells positively staining for LAG-3 and TIGIT (**Fig. 2B**).

**Figure 2.**
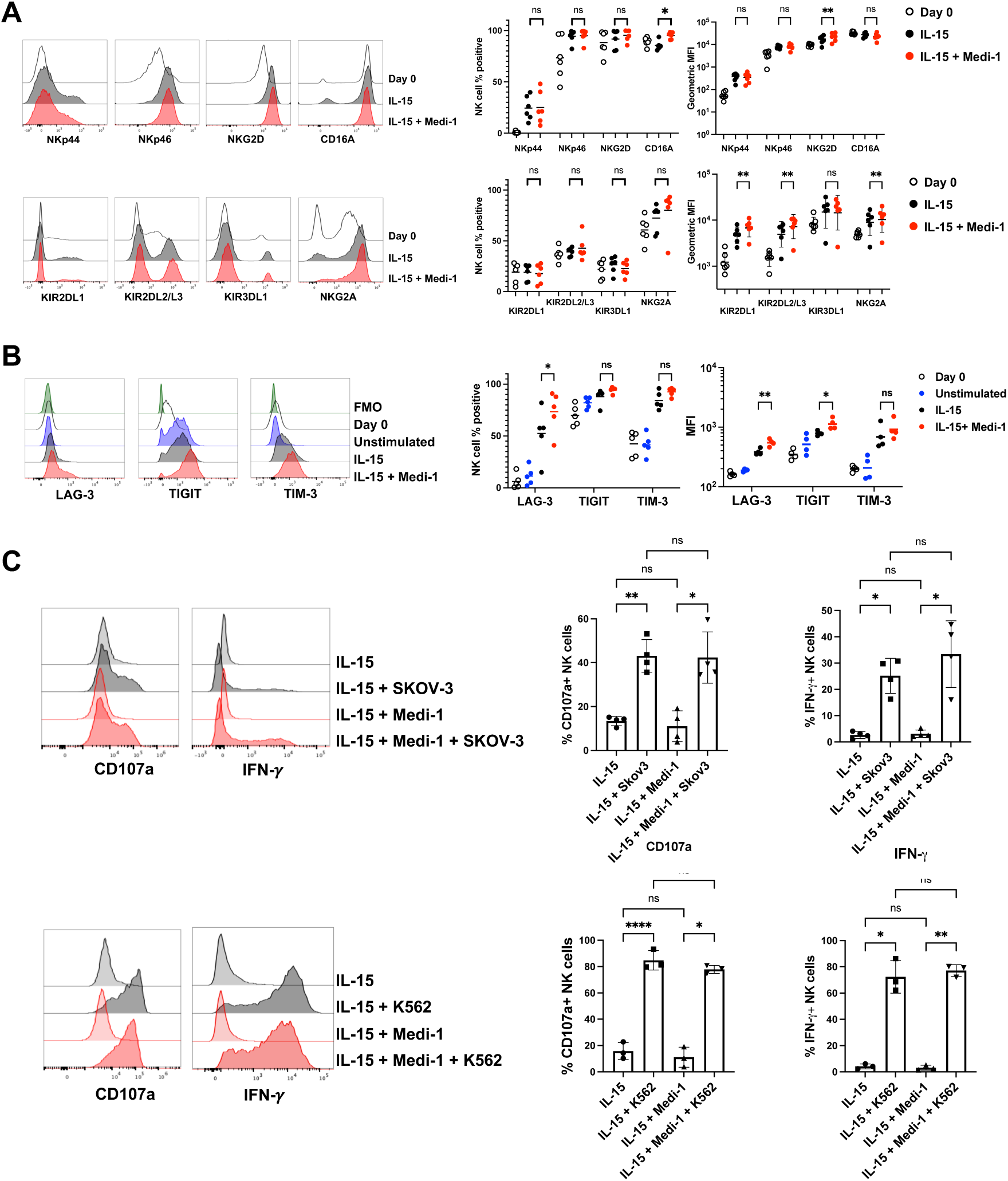
Proliferating NK cells treated with IL-15 +/- Medi-1 express similar functional markers. Freshly isolated PBMCs were cultured in the presence or absence of IL-15 (10ng/ml) and Medi-1 (5µg/ml), as indicated. At day 0 or day 7, CD56^+^ CD3^−^ NK cells were analyzed for their expression levels of the indicated markers by flow cytometry. **A.** Activating receptors (top panels) and inhibitory receptors (bottom panels). Representative histogram plots (left panels). y-axis = cell number. Cumulative data showing the percentage of NK cells positively staining for each marker (mean is indicated by the horizontal line) and their geometric MFI (right panels). For the latter, data were log-transformed prior to analysis. n=6 donors. **B.** Exhaustion markers. Representative histograms are shown in the left panel. Unstimulated = unstimulated cells stained at day 7. Fluorescence minus one (FMO) of cells at day 0. Cumulative data are shown as the percentage of NK cells positively staining for each marker and the geometric MFI of staining. For the latter, data were log-transformed prior to analysis. n=4-5 donors. **C.** PBMCs were cultured in IL-15 +/- Medi-1 for 7 days, washed, and then incubated with K562 or SKOV-3 cells for 5 hours (effector: target ratio = 1:1). PBMCs were stained for CD56, CD3, CD107a, and IFN-γ and examined by flow cytometry (CD56^+^ CD3^−^ cells are shown). Representative histogram plots (left panels) and cumulative data (right panels) are shown, n=3-4 donors. *p < 0.05; **p < 0.01; ****p < 0.0001; ns=not significant. Statistical significance was determined by a one-way ANOVA with a Dunnett post hoc test.

To assess the significance of the modest increase in the expression of certain activating and inhibitory receptors by NK cells treated with IL-15 and Medi-1, we evaluated the functional readouts of degranulation and cytokine production via CD107a and IFN-γ detection, respectively, which are diminished upon NK cell exhaustion ^23^. PBMCs were stimulated with IL15 +/- Medi-1 for 7 days and then co-cultured with the ovarian cancer cell line SKOV-3 or the myelogenous leukemia cell line K562 for 5 hours. With both tumor cell targets, NK cells expanded by IL-15 +/-Medi-1 upregulated CD107a and IFN-γ at equivalent levels (**Fig. 2C**), though their response to K562 cells was much more robust. Hence, though Medi-1 treatment augmented IL-15-driven NK cell proliferation and increased the expression of certain inhibitory markers, this did not lead to broad changes in well-recognized functional phenotypic markers.

### Early activation markers distinguish IL-15-stimulated NK cells treated with Medi-1

We next examined the expression of the early activation markers CD69, CD25 (IL-2Rα), and CD137 (4-1BB) on NK cells following the stimulation of PBMCs with IL-15 +/- Medi-1. Upon IL-15 stimulation, NK cells rapidly and uniformly upregulated CD69, which remained elevated for the 7-day culture (**Fig. 3**). IL-15 stimulation along with Medi-1 increased CD69 upregulation on day 1, but its expression modestly decreased at the remaining time points (**Fig. 3**). CD25 levels on NK cells following IL-15 stimulation were increased at all time points compared to unstimulated NK cells, and this was markedly enhanced in the presence of Medi-1 on days 1–4 (**Fig. 3**). Similarly, Medi-1 treatment significantly enhanced CD137 upregulation on NK cells by IL-15 (**Fig. 3**). Hence, CD25 and CD137, but not CD69, demonstrated distinct expression patterns on NK cells treated with IL-15 and Medi-1 when compared to IL-15 treatment alone, which was apparent by day 1 of stimulation (**Fig. 3**).

**Figure 3.**
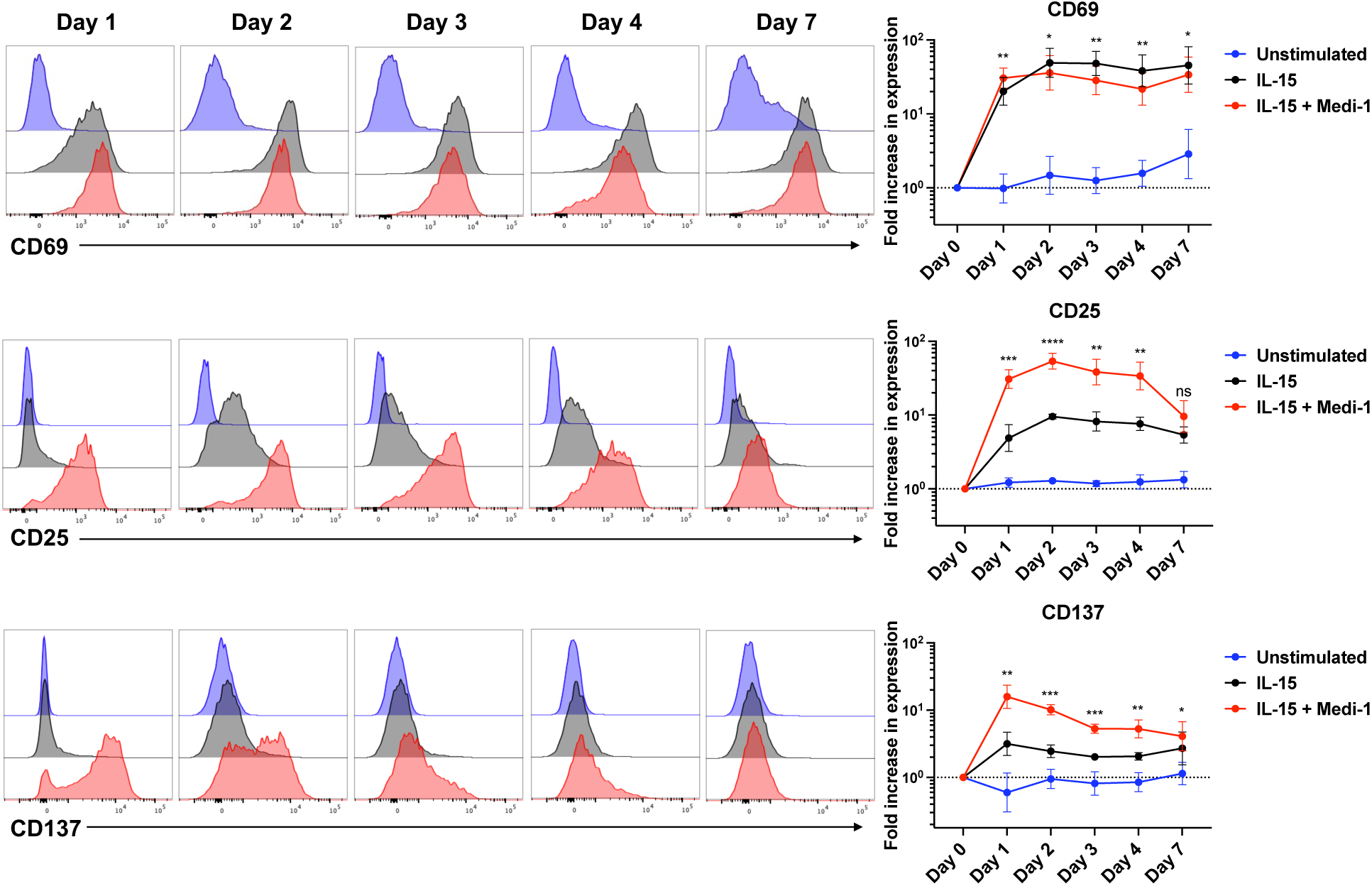
Medi-1 treatment augments the upregulation of early activation markers on NK cells. Freshly isolated PBMCs were treated as described in Fig. 2. CD56^+^ CD3^−^ NK cells were analyzed for their CD69, CD25, and CD137 expression levels by flow cytometry on days 0, 1-4, and 7 of culture. Representative histogram plots (left panels). y-axis=cell number. Cumulative data (right panels) are shown as an averaged log transformed geometric MFI fold increase from day 0. Mean +/- SD, n=4 donors. *p < 0.05; **p < 0.01; ***p < 0.001; ****p < 0.0001; ns=not significant. Statistical significance was determined by a two-way ANOVA with Tukey post hoc test.

The kinetics of CD69, CD25, and CD137 upregulation were also examined using enriched NK cells treated with IL-15 +/- Medi-1 for up to 7 days. Again, CD25 and CD137 demonstrated a distinct pattern of upregulation by IL-15-stimulated NK cells treated with Medi-1 (**Fig. 4A**). Since CD137 achieved peak expression by day 1, we examined earlier time points of stimulation at 4, 8, and 18 hours. CD25 and CD69 demonstrated significantly higher levels of upregulation as early as 4 hours after NK cell treatment with IL-15 and Medi-1 compared to NK cells stimulated with IL-15 alone (**Fig. 4B**). Medi-1 treatment significantly increased CD137 as early as 8 hours after stimulation relative to NK cells treated with IL-15 alone (**Fig. 4B**). These data demonstrate that Medi-1 treatment played a direct role in enhancing NK cell activation by IL-15 (i.e., independent of other PBMC populations).

**Figure 4.**
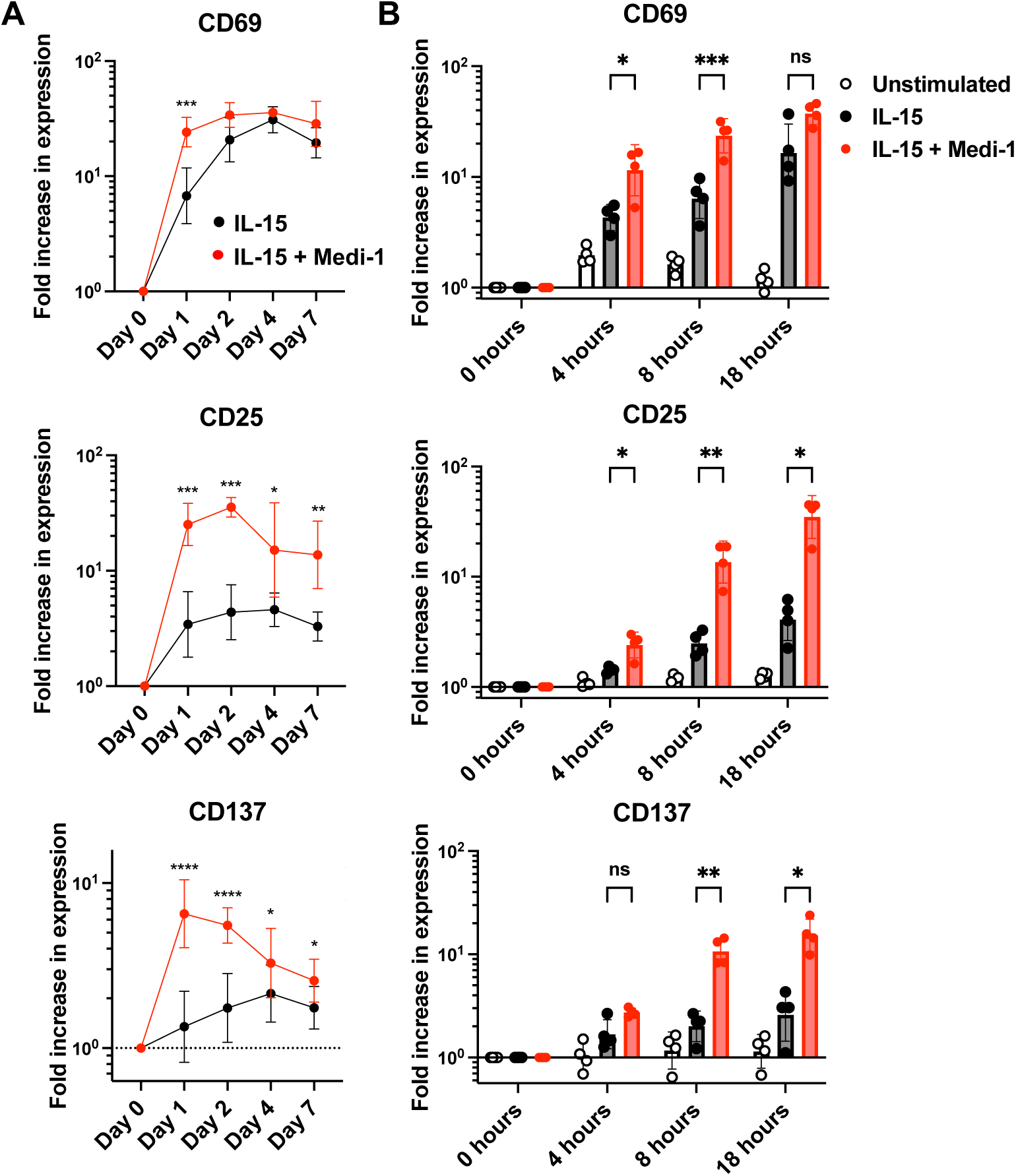
Direct effects of Medi-1 on enriched NK cells. Freshly isolated NK cells (>90% enriched) were treated and analyzed as described in Fig. 2. **A.** CD56^+^ CD3^−^ NK cells were analyzed for their expression levels of CD69, CD25, and CD137 by flow cytometry on days 0, 1-4, and 7 of culture. **B.** CD56^+^ CD3^−^ NK cells were similarly examined at hours 0, 4, 8, and 18. Data are shown as the averaged log-transformed geometric MFI fold increase from day 0 (A) hour 0 (B). Mean +/- SD, n=4 donors. *p < 0.05; **p < 0.01; ***p < 0.001; ****p < 0.0001. Statistical significance was determined by a two-way ANOVA with a Šídák (A) or Tukey post hoc test (B).

### The Fc region of Medi-1 enhances NK cell activation by IL-15

The distinct expression patterns of CD25 and CD137 by IL-15-stimulated NK cells when treated with Medi-1 were indicative of its synergetic effect. CD16A is the main IgG Fc activating receptor expressed by human NK cells and it exclusively mediates ADCC ^10^. As Medi-1 is a human IgG1 mAb, it can potentially be engaged by CD16A upon binding NK cells. To test whether the Fc region of Medi-1 contributed to its synergistic effect, we generated several other Fc variants of the mAb, including Medi-F(ab′)_2_ in which its Fc region was removed; Medi-PGLALA containing the mutations Pro329Gly, Leu234Ala, and Leu235Ala that also abolish FcγR binding, but with less potential effects on antibody structure and function ^25^; and Medi-4 containing the Fc region of IgG4, which binds to CD16A with ∼10-fold lower affinity than IgG1^26^. Hence, the rank order of CD16A binding affinity for the generated Medi Fc variants is Medi-1>Medi-4> Medi-PGLALA>/= Medi-F(ab′)_2_. All versions of Medi, but not a human IgG control mAb, equivalently blocked CD16A downregulation upon cell activation (**Supplemental Fig. 2**), demonstrating that their ADAM17 blocking function was not altered. PBMCs were stimulated with IL-15 +/- a Medi Fc variant at the same molar concentration for 24 hours. The expression of CD25 and CD137 was then determined by flow cytometry. As shown in **Figure 5A**, IL-15-stimulated NK cells treated with Medi-1 vs. the other Fc variants demonstrated significantly higher levels of these activation markers. The same effects were observed on IL-15-driven NK cell proliferation (**Fig. 5B**).

**Figure 5.**
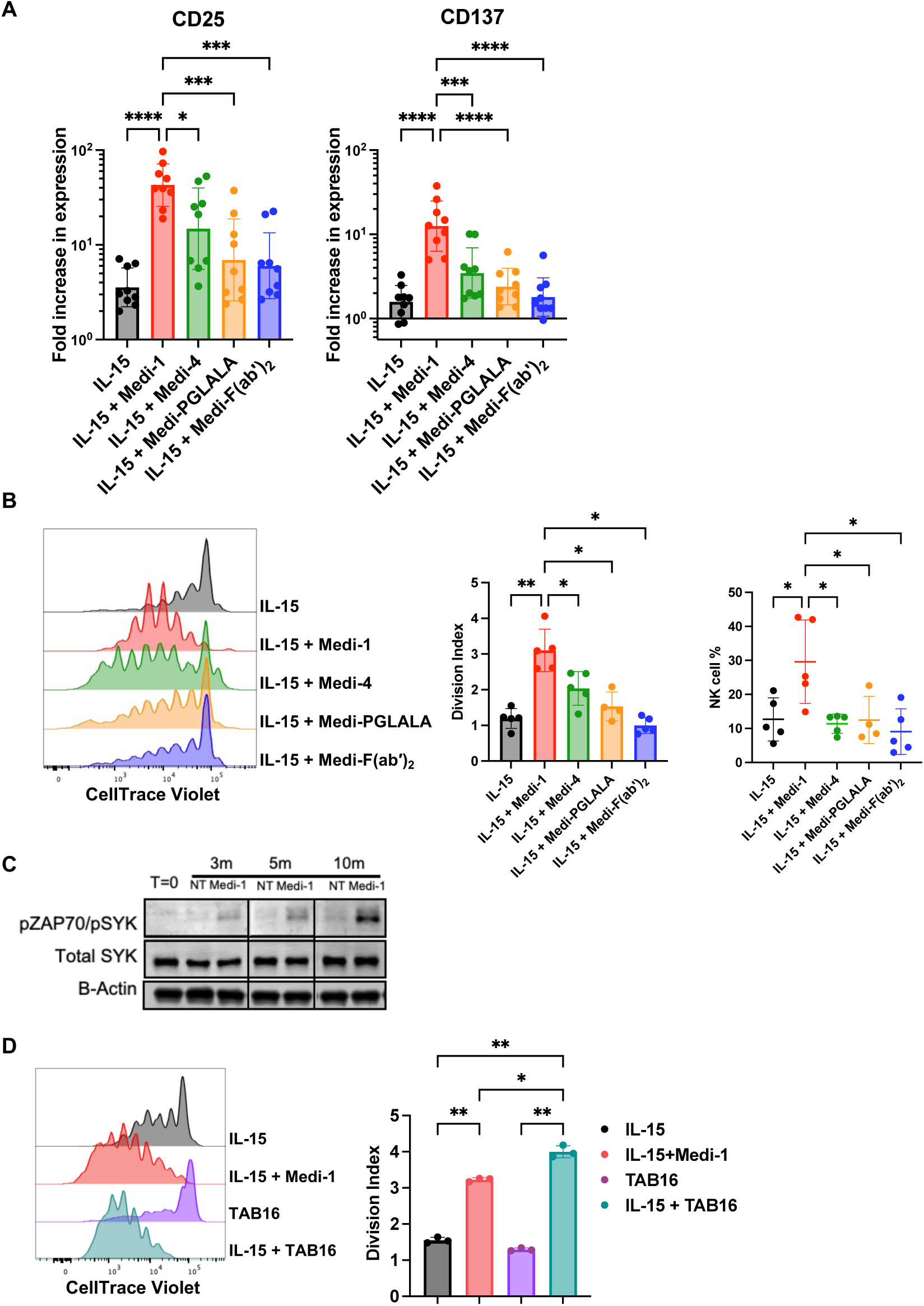
Medi-1 synergizes with IL-15 stimulation through its Fc region. **A.** Freshly isolated PBMCs were cultured with IL-15 (10ng/ml) +/- molar equivalents of Medi-1, Medi-4, Medi- PGLALA, or Medi-F(ab′)_2_, as indicated. CD56^+^ CD3^−^ NK cells were analyzed for their expression levels of CD25 and CD137 by flow cytometry at 18 hours of culture. Data are shown as the averaged log-transformed geometric MFI fold increase from day 0. Mean +/- SD, n=9 donors. **B.** PBMCs were labeled with CTV and cultured for 7 days with IL-15 +/- Medi-1, Medi-4, Medi-PGLALA, or Medi-F(ab′)_2_, as indicated. CD56^+^ CD3^−^ NK cells were analyzed for CTV dilution by flow cytometry. Representative histograms (left panel) and division index (middle panel) are shown. Percentage of NK cells for the indicated conditions (right panel). Mean +/- SD, n=5 donors. **C.** Enriched NK cells were cultured with Medi-1 or no antibody (no treatment, NT) for the indicated times. Immunoblotting shows pZap-70/pSyk and total Syk. β-actin indicates protein loading. Representative data are shown. **D.** PBMCs were treated as described in panel B with IL-15 +/- Medi-1 or TAB16 at equivalent molarity (33nM). Representative histograms (left panel) and division index (right panel) are shown. Mean +/- SD, n=3 donors. *p < 0.05; **p < 0.01; ***p < 0.001; ****p < 0.0001. Statistical significance was determined by a one-way ANOVA with a Dunnett (A, D) or Tukey (B) post hoc test.

The above findings indicate that CD16A engages the Fc region of Medi-1 when attached to NK cells. CD16A non-covalently associates with the signaling adaptors FcRγ and CD3ζ ^27 28^. These immunoreceptor tyrosine-based activation motif (ITAM)-containing signaling adaptors become phosphorylated and associate with the Syk family protein tyrosine kinase Zap-70, which is also phosphorylated, leading to downstream signaling cascades for cell activation that include PLCγ1 and ERK phosphorylation ^29^. We observed that treating NK cells with Medi-1 alone led to detectable levels of Zap-70 phosphorylation (**Fig. 5C**).

As an approach to directly engage CD16A and simultaneously block ADAM17 function, we generated a bispecific antibody engager that consists of a humanized camelid nanobody against CD16 and a scFv from Medi, referred to as a Targeted ADAM17 Blocker-16 (TAB16). PBMCs were labeled with CTV and cultured with IL-15 alone or with molar equivalents of Medi-1 or TAB16 for 7 days. TAB16 + IL-15 induced significantly higher levels of NK cell proliferation than IL-15 alone, and, interestingly, higher levels of proliferation than IL-15 + Medi-1 (**Fig. 5D**).

A potential adverse effect of CD16A engagement of the Fc region Medi-1 would be the induction of NK cell degranulation and fratricide. For example, daratumumab, a human IgG1 mAb that recognizes CD38 on NK cells, has been shown to induce fratricide ^30^. Indeed, we observed impaired NK cell proliferation and a greatly decreased representation within the PBMC population when CTV-labelled PBMCs were cultured for 7 days with IL-15 and daratumumab, which was not seen with Medi-1 treatment (**Fig. 6A**), indicating it did not induce a critical level of fratricide or clearance by macrophage present in the culture. In other assays, enriched NK cells cultured with daratumumab for 24 hours demonstrated a significant upregulation in cell surface levels of CD107a, a sensitive marker of NK cell degranulation upon CD16A signaling ^8^, compared to control cells. This was not observed for NK cells cultured with Medi-1 (**Fig. 6B**). To assess the effects of Medi-1 treatment on the functional state of NK cells, we examined their natural cytotoxicity against tumor cells. For this assay, enriched NK cells were used to exclude anti-tumor activity by other leukocytes. NK cells were cultured with SKOV-3/NLG cells at various E:T ratios in the presence of IL-15 +/- Medi-1, and tumor cell lysis was assessed by IncuCyte live cell imaging for 7 days. Interestingly, in the presence of Medi-1, NK cells mediated significantly higher levels of SKOV-3 killing (**Fig. 6C**). Taken together, our results show that the Fc region of Medi-1 contributed to the activation and proliferation of IL-15-stimulated NK cells while not inducing overt adverse effects.

**Figure 6.**
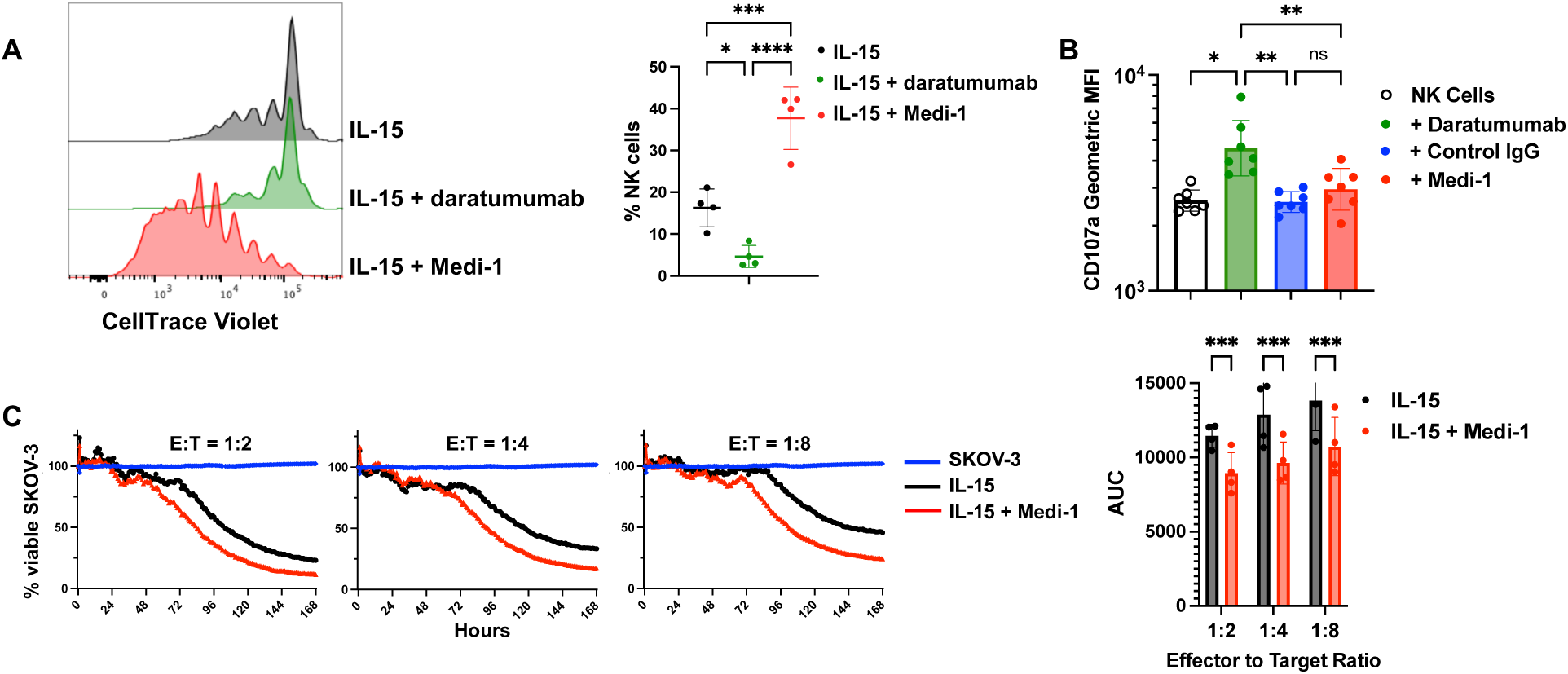
Medi-1 treatment of IL-15-stimulated NK cells does not induce fratricide or impair natural cytotoxicity. **A.** PBMCs were labeled with CTV and cultured for 7 days with IL-15 (10ng/ml) +/- Medi-1 or daratumumab at 5µg/ml, as indicated. CD56^+^ CD3^−^ NK cells were analyzed for CTV dilution by flow cytometry. Representative data are shown (left panel) as well as the percentage of NK cells for the indicated conditions (right panel). Mean +/- SD, n=4 donors. **B.** PBMCs cells were cultured for 24 hours in IL-15 +/- an isotype-matched negative control mAb (human IgG1), Medi-1 (5µg/ml), or daratumumab at (5µg/ml), as indicated. CD56^+^ CD3^−^ NK cells were analyzed for their expression levels of CD107a by flow cytometry. Cumulative data are shown as geometric MFI +/- SD, n=7 donors. **C.** Enriched NK cells were co-cultured with SKOV-3-NLG cells at the indicated E:T ratios for 7 days in the presence of IL-15 (10ng/ml) +/- Medi-1 (5µg/ml). Cytotoxicity was assessed by IncuCyte-based live cell imaging. The percentage of viable SKOV-3 cells was double normalized to SKOV-3 cells alone (left panel). The area under the curve (AUC) of the remaining SKOV-3 was calculated for each condition. Mean +/- SD, n=4. *p < 0.05; **p < 0.01; ***p < 0.001. Statistical significance was determined by a one-way ANOVA with Tukey post hoc test (A, B) or two-way ANOVA with Šídák post hoc test (C).

### Enhanced IL-15-driven NK cell proliferation by Medi-1 involves CD137 and PBMC accessory cells

Circulating NK cells are composed of CD56 dim and bright populations, and CD16A is primarily expressed by CD56^dim^ NK cells ^31^. It was mainly these cells that upregulated CD137 after stimulation with Medi-1 and IL-15 (**Supplemental Fig. 4**). It is well established that CD137 stimulation in T cells promotes their activation and proliferation ^32^. However, much less is known about the biological importance of CD137 in NK cells. Engagement of this co-activation receptor by agonistic antibodies or by CD137L-expressing artificial feeder cells induces NK cell proliferation ^33 34^. To investigate the role of CD137 in the augmented proliferation of Medi-1 + IL-15-stimulated NK cells, we blocked its function. Others have reported that the addition of soluble CD137L to PBMC cultures can inhibit T cell proliferation by acting as a competitive inhibitor ^35 36^. We attempted this for NK cells using a recombinant human CD137L extracellular domain-Fc fusion protein (sCD137L-Fc). We found that sCD137L-Fc diminished NK cell proliferation by IL-15 as well as IL-15 + Medi-1 (**Fig. 7A**), which occurred in a dose-dependent manner (**Supplemental Fig. 5**). A caveat of this approach is that sCD137L-Fc, when attached to NK cells, might be engaged by CD16A and reduce its engagement of Medi-1. However, the presence of sCD137L-Fc did not affect the enhanced upregulation of CD25 by NK cells treated with IL-15 and Medi-1 (**Fig. 7B**), indicating that Medi-1 engagement by CD16A was not affected. We could not assess CD137 staining since sCD137L-Fc competed with our anti-CD137 mAb used for flow cytometry (data not shown), demonstrating its attachment to CD137.

**Figure 7.**
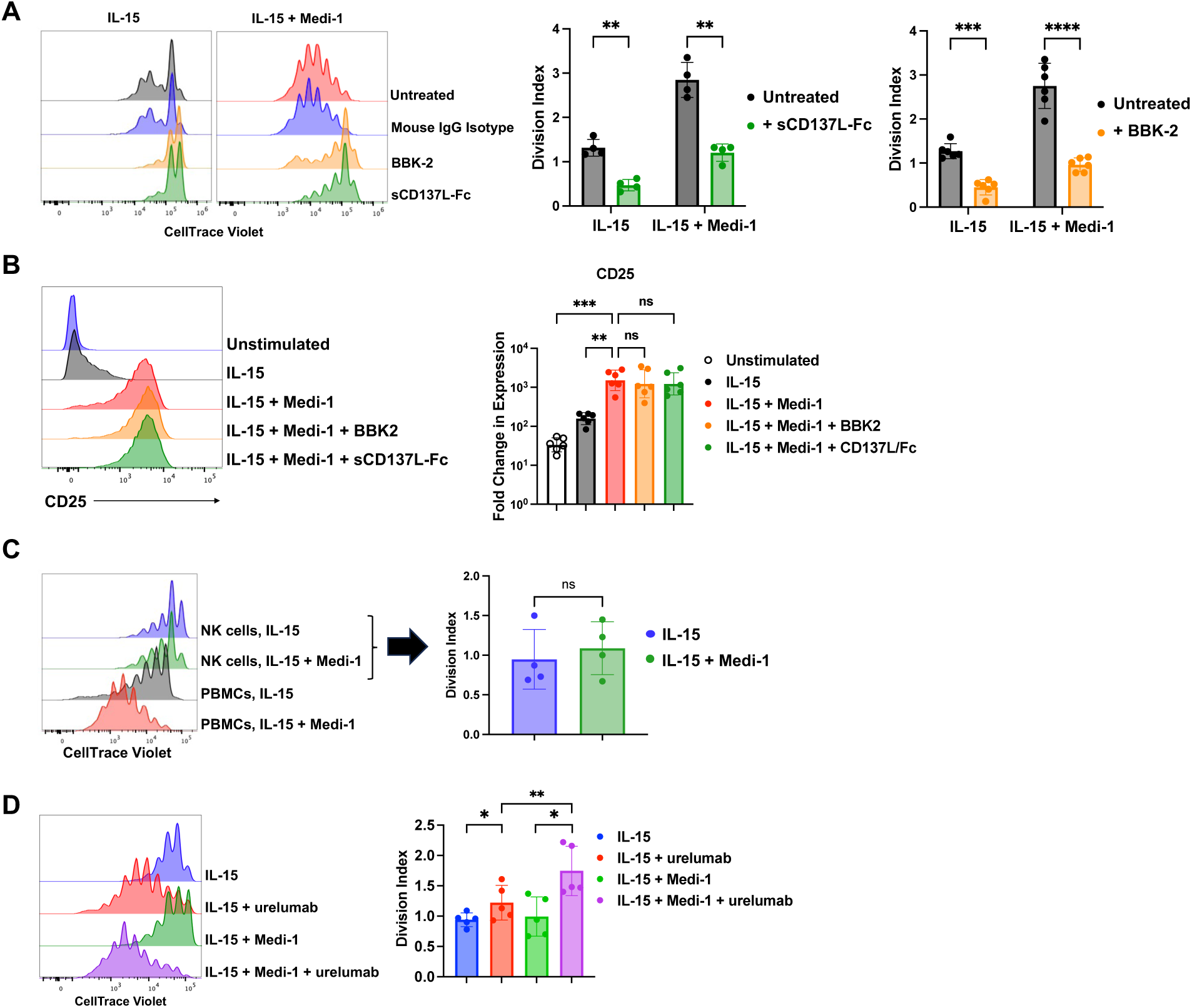
Medi-1-enhanced NK cell proliferation by IL-15 involves CD137. PBMCs were labeled with CTV and cultured for 7 days with IL-15 (10ng) +/- Medi-1 (5µg/ml) +/- BBK-2 (5µg/ml), an isotype-matched negative control mAb (mouse IgG1, 5µg/ml), or sCD137L-Fc fusion protein (1µg/ml). **A.** CD56^+^ CD3^−^ NK cells were analyzed for CTV dilution by flow cytometry (left panel). Division index of NK cells in the presence or absence of BBK-2 or sCD137L-Fc fusion protein (right panels). Mean +/- SD, n=4-6 donors. **B.** PBMCs were treated with IL-15 +/- the indicated reagents for 24 hours and the expression of CD25 was determined on NK cells. Representative CD25 expression on NK cells (left panel). The right panel shows the log-transformed fold change in CD25 geometric MFI from baseline. Mean +/- SD, n=6. **C.** CTV-labeled PBMCs or enriched NK cells from matched donors were cultured for 7 days with IL-15 (10ng) +/- Medi-1 (5µg/ml). CD56^+^ CD3^−^ NK cells were analyzed for CTV dilution by flow cytometry (left panel). Division index (right panel) for enriched NK cells, mean +/- SD. n=4 donors. **D.** CTV-labeled enriched NK cells were cultured for 7 days with IL-15 (10ng) +/- Medi- 1 (5µg/ml) +/- urelumab (1µg/ml). CD56^+^ CD3^−^ NK cells were analyzed for CTV dilution by flow cytometry (left panel). Division index (right panel), mean +/- SD. n=5 donors. *p < 0.05; **p < 0.01; ***p < 0.001; ****p < 0.0001. Statistical significance was determined by a two-way ANOVA with Šídák post hoc test (A), one-way with Dunnentt’s post hoc test (B) or paired two-tailed Student’s t-tests (C, D).

We also examined the effects of a function-blocking anti-human CD137 mAb on NK cell proliferation by IL-15. Anti-CD137 mAbs that inhibit its attachment to CD137L can also be agonistic and induce cell activation, as is the case for utomilumab ^37^. To our knowledge, the best characterized blocking anti-human CD137 mAb that is not agonistic is BBK-2, which partially neutralizes CD137 function ^38^. PBMCs were stimulated with IL-15 +/- Medi-1 in the presence or absence of BBK-2 or an appropriate isotype control mAb. BBK-2 and the control mAb obtained from commercial sources contained NaN_3_ as a preservative. We have found that even very low concentrations of NaN_3_ can impair NK cell activation and proliferation (data not shown). Therefore, to prevent a confounding effect by the preservative, it was removed from both commercially sourced antibodies by a desalting procedure, as described in the Material and Methods. As shown in **Figure 7A**, BBK-2, but not the isotype control mAb, significantly reduced NK cell proliferation by IL-15 alone and in combination with Medi-1. Taken together, the above findings support CD137 upregulation as a mechanism by which Medi-1 treatment augments IL-15-mediated NK cell proliferation. CD137L is primarily expressed by monocytes, macrophages, and B cells, and it can be upregulated following their activation ^39^. Interestingly, we observed that only IL-15-stimulated NK cells in PBMCs and not enriched NK cells (≥ 90% NK cell purity) underwent enhanced proliferation in the presence of Medi-1 (**Fig. 7C**), demonstrating an important role for PBMC accessory cells.

To address whether CD137 activation could enhance NK cell proliferation by IL-15 in the absence of PBMC accessory cells, we used an anti-CD137 agonist mAb. Enriched NK cells were labeled with CTV and cultured for 7 days with IL-15 +/- Medi-1 +/- urelumab, a well-described CD137 agonistic mAb ^40^. Interestingly, in the presence of urelumab, enriched NK cells underwent a marked increase in proliferation by IL-15, and this was increased further by the addition of Medi-1 (**Fig. 7D**).

## DISCUSSION

We show that the ADAM17 function-blocking mAb Medi-1 markedly enhanced NK cell activation and proliferation by IL-15, and that the latter required the presence of PBMC accessory cells. In this study, we investigated the mechanisms by which Medi-1 augmented these events. IL-15 stimulated NK cells treated with Medi-1, but not IL-15 or Medi-1 alone, distinctly upregulated high levels of CD25 and CD137, indicating a synergistic activation process. Medi-1 is a human IgG1 antibody and when bound to NK cells it could be engaged by their activating IgG Fc receptor CD16A. Indeed, NK cell treatment with various Medi Fc variants in which the Fc region was removed (Medi-F(ab’)_2_), mutated (Medi-PGLALA), or exchanged with the Fc region of IgG4 to prevent or diminish CD16A binding resulted in significantly reduced NK cell activation and proliferation in the presence of IL-15 when compared to Medi-1. In addition, we show that CD16A signaling was induced when NK cells were treated with Medi-1. To directly co-engage ADAM17 and CD16A, we generated the unique bi-specific antibody TAB16 consisting of an scFv from Medi-1 and a camelid antibody specific to CD16. TAB16 treatment also significantly enhanced IL-15-driven NK cell proliferation, more so than Medi-1 and IL-15. This could be due to a higher affinity interaction between the anti-CD16 camelid antibody and CD16A than CD16A binding to the Fc region of Medi-1, in turn inducing increased CD16A signaling. Indeed, signaling by CD16A when binding to NK cell-attached Medi-1 will likely vary due to CD16A polymorphisms distributed within the normal population that affect its binding affinity ^41^.

An issue with CD16A engagement of NK cell bound antibodies is the induction of fratricide, which has been reported for the anti-CD38 mAb daratumumab ^30^. We show, however, that Medi-1 treatment did not induce degranulation by enriched NK cells nor their elimination in PBMC cultures, whereas daratumumab treatment caused both events. CD16A is a low affinity FcγR and its signaling upon binding antibody-opsonized cells is affected by antigen density, mobility, and accessibility. For instance, low antigen density is ADCC limiting, as a threshold of CD16A signaling is required to induce degranulation by NK cells ^21^. ADAM17 can be constitutively expressed at low levels on the surface of cells ^42^, including hematopoietic cells ^43^, and so the surface density of ADAM17 on NK cells may be low enough that Medi-1 engagement by CD16A does not achieve the signaling threshold necessary for degranulation in a critical mass of cells. NK cell effector activities induced by CD16A may vary depending on its signaling level, akin to T cell effector functions modulated by TCR signaling strength ^44^.

Despite the enhanced activation and proliferation of IL-15-stimulated NK cells by Medi-1, we did not observe overt changes in their expression of several functional markers when compared to NK cells stimulated with IL-15 alone that would indicate exhaustion or dysfunction. To the contrary, we observed that NK cells stimulated with IL-15 in the presence of Medi-1 mediated significantly higher levels of tumor cell natural cytotoxicity at various E:T ratios. Currently, we do not know the cause of this increased killing. It is possible that CD16A signaling induced by Medi-1 synergized with other activating receptors on NK cells to enhance their cytolytic activity. Addressing this and the effects of Medi-1 on the anti-tumor activity of NK cells *in vivo* are important future studies. However, the latter would likely be due to more than just contributions by CD16A considering that ADAM17 has several substrates ^11^.

D1(A12), another human IgG1 ADAM17 function-blocking mAb ^20^, did not enhance NK cell activation and proliferation by IL-15. There are a few explanations that may account for the different effects of D1(A12) and Medi-1 on NK cells, including epitope differences, binding affinity, and Fc glycosylation and/or polymorphisms that could affect CD16A engagement ^21^. It was beyond the scope of our study to determine the underlying reason for this difference. Our findings show, however, that not all ADAM17 function-blocking mAbs may be equivalent in enhancing IL-15-driven NK cell proliferation.

Of interest was the robust upregulation of CD137 by NK cells treated with IL-15 and Medi-1. CD137 is a potent co-activating receptor that has mainly been studied in T cells. CD8 T cells undergoing antigen activation and CD137 engagement demonstrate enhanced proliferation, reversal of exhaustion, apoptosis protection, and increased cytokine secretion and other effector functions ^32^. In contrast, the biological function of CD137 in NK cells is not well understood. IL-15 stimulation and CD16A engagement both induce CD137 upregulation in NK cells ^34 45^. Nuclear factor κB (NF-κB) and nuclear factor of activated T cells (NFAT) are involved in the transcriptional induction of CD137 ^46 47^, and they are activated by IL-15 and CD16A signaling, respectively ^46 48^. Though the molecular mechanisms underlying the effects of Medi-1 and IL-15 on CD137 expression by NK cells are not currently known, it is possible that NF-κB and NFAT work in synergy to drive this process.

PBMC accessory cells are known to enhance NK cell proliferation by transpresenting IL-15 ^18 49^. CD137L is primarily expressed by monocytes, macrophages, and B cells, and it can be upregulated following their activation ^39^. We demonstrate for the first time that NK cell proliferation by IL-15 in the presence of PBMC accessory cells can be diminished by blocking CD137, as was their enhanced proliferation mediated by Medi-1. CD137 upregulation by IL-15-stimulated NK cells may establish a positive feedback loop to increase NK cell proliferation, which can be further augmented by Medi-1 treatment. Of interest is that the induction of CD137 is targetable by the agonistic CD137 antibody urelumab. This approach to amplify NK cell expansion could have applications in patients and *ex vivo* prior to their adoptive transfer as a feeder cell-free method.

In summary, we have demonstrated that Medi-1 treatment synergizes with IL-15 in activating NK cells and enhancing their proliferation. IL-2 is also broadly used for NK cell stimulation and proliferation *ex vivo* and in cancer patients ^7^, and we show that Medi-1 treatment of NK cells similarly enhanced NK cell proliferation by this cytokine. Thus, the synergistic activity of Medi-1 on NK cells was not specific to IL-15. The mechanisms underpinning Medi-1’s effects involve its engagement by CD16A and blocking ADAM17, which induce and presumably prolong its signaling. This increased the upregulation of CD137 expression, promoting NK cell expansion in the presence of PBMC accessory cells expressing CD137L (**Graphical abstract**). Our study is of translational importance as Medi-1 treatment in combination with cytokine therapies could potentially augment the proliferation and function of endogenous or adoptively transferred NK cells in cancer patients.

## Abbreviations

NK cell: natural killer cell
PBMC: peripheral blood mononuclear cell
TriKE: Trispecific Killer Engager
CTV: CellTrace Violet
ADAM17: a disintegrin and metalloproteinase-17
ADCC: antibody-dependent cell-mediated cytotoxicity
NaN_3_: sodium azide
FMO: fluorescence minus one
E:T: effector:target
PMA: phorbol 12-myristate 13-acetate
MFI: mean fluorescent intensity
AUC: Area under the curve
TAB16: Targeted ADAM17 Blocker-16

## SUPPLEMENTAL FIGURE LEGENDS

**Supplemental Figure 1.**
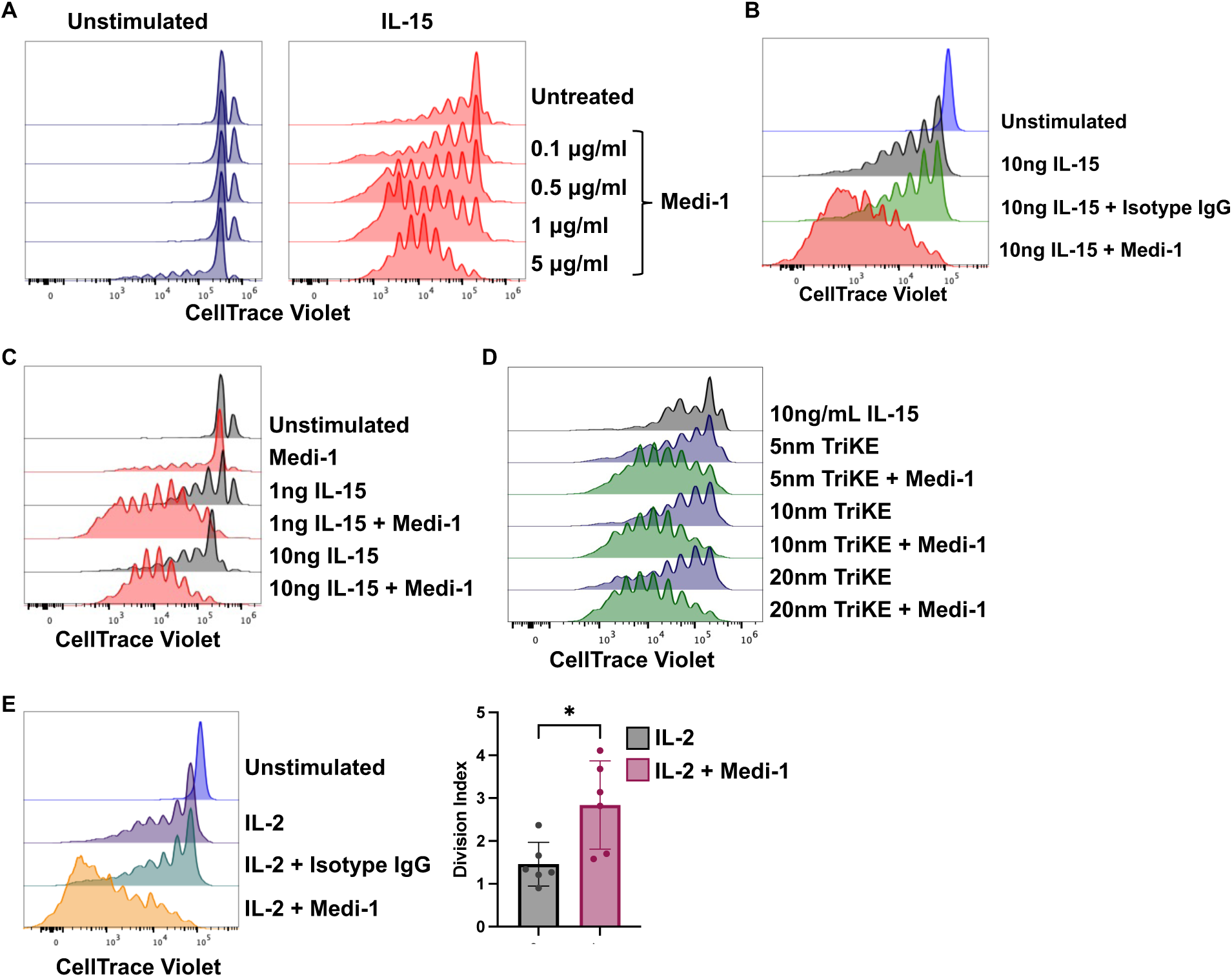
Enhanced IL-15-driven NK cell proliferation by Medi-1. **A, B.** Freshly isolated PBMCs were CTV-labeled and cultured for 7 days +/- IL-15 (10ng/ml) +/- Medi-1 at the indicated concentrations (A) or an isotype-matched negative control mAb (human IgG1) (5 µg/ml) (B). **C.** CTV-labeled PBMCs were cultured for 7 days +/- IL-15 at the indicated concentrations +/- Medi-1 (5 µg/ml). **D.** CTV-labeled PBMCs were cultured for 7 days +/- cam16-IL15-camB7H3 (TriKE) at the indicated concentrations +/- Medi-1 (5 µg/ml). CD56^+^ CD3^−^ NK cells were analyzed for CTV dilution by flow cytometry. Data are representative of 3 independent experiments using leukocytes from separate donors. **E.** CTV-labeled PBMCs were cultured for 7 days with IL-2 (200 IU/ml) +/- Medi-1 (5 µg/ml) or an isotype-matched negative control mAb (human IgG1) (5 µg/ml). CD56^+^ CD3^−^ NK cells were analyzed for CTV dilution by flow cytometry. Representative data are shown (left panel). Division index (right panel), mean +/- SD, n=6 donors. *p < 0.05; ns=not significant. Statistical significance was determined by paired two-tailed Student’s t-tests.

**Supplemental Figure 2.**
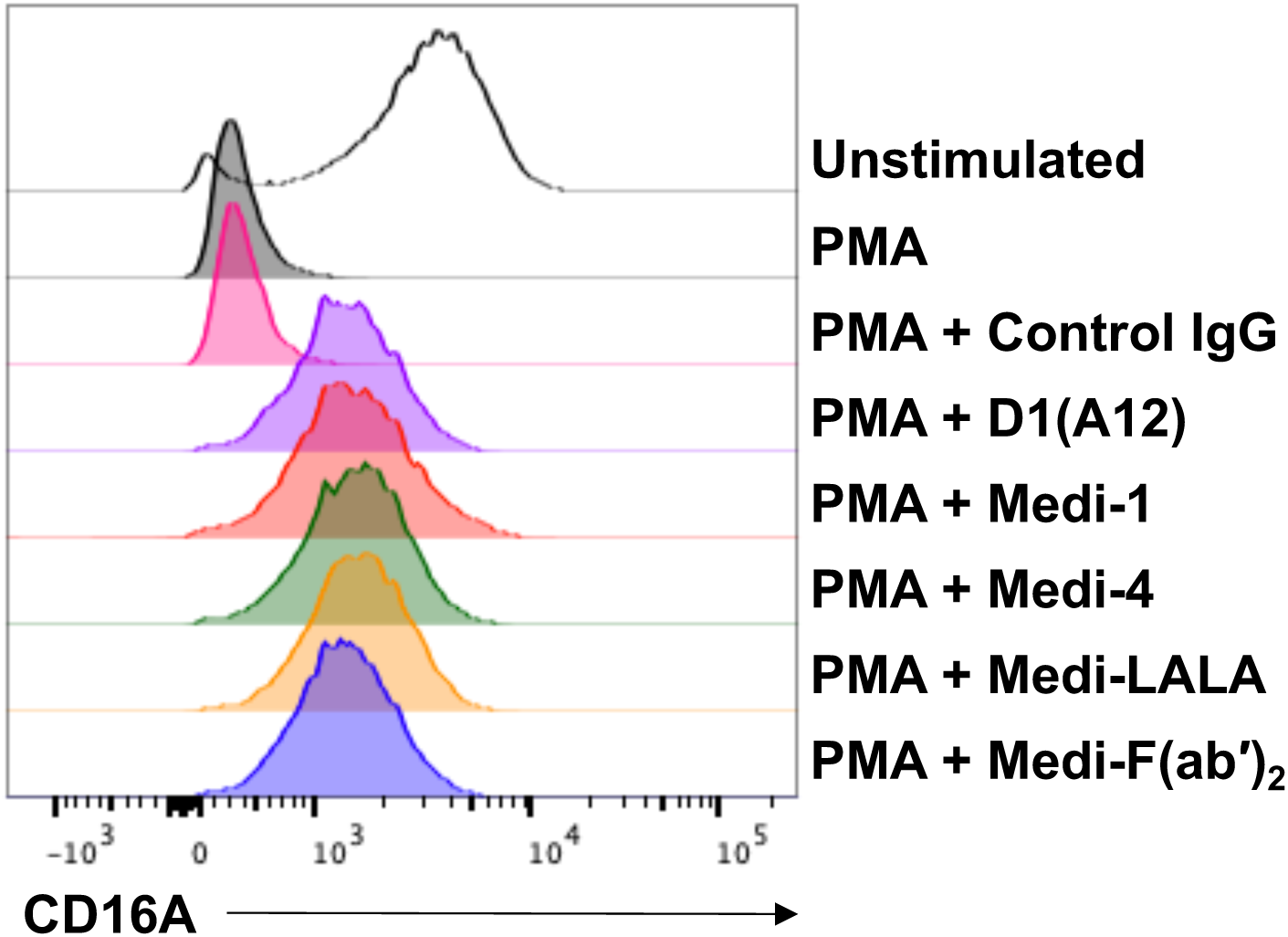
Medi-Fc variants block ADAM17 function. NK-92 cells expressing CD16A were treated with or without PMA/ionomycin in the presence or absence of molar equivalents (33 nM) of an isotype-matched negative control mAb (human IgG1), Medi-1, Medi-4, Medi-PGLALA, or Medi-F(ab′)_2_. Relative cell-staining levels of CD16A were determined by flow cytometry. Data are representative of 3 independent experiments.

**Supplemental Figure 3.**
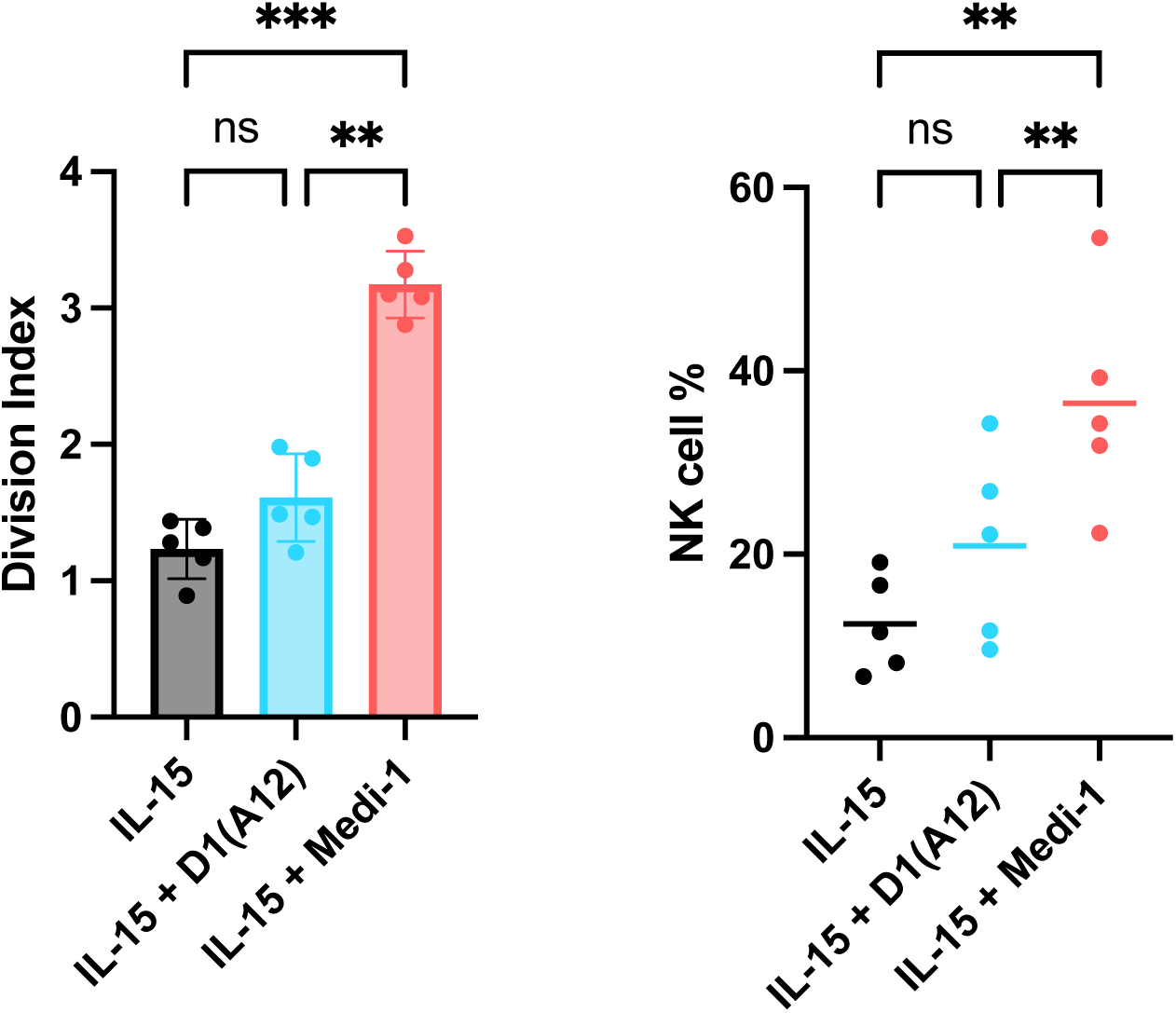
The function-blocking anti-ADAM17 mAb D1(A12) did not enhance IL-15-driven NK cell proliferation. Freshly isolated PBMCs were CTV-labeled and cultured for 7 days with IL-15 (10 ng/ml) +/- D1(A12) (5 μg/ml) or Medi-1 (5 μg/ml). CD56^+^ CD3^−^ NK cells were analyzed for CTV dilution by flow cytometry. Division index (left panel) and percentage of NK cells for the indicated conditions (right panel). **p < 0.01; ***p < 0.001; ns=not significant. Statistical significance was determined by a one-way ANOVA with Tukey post hoc test.

**Supplemental Figure 4.**
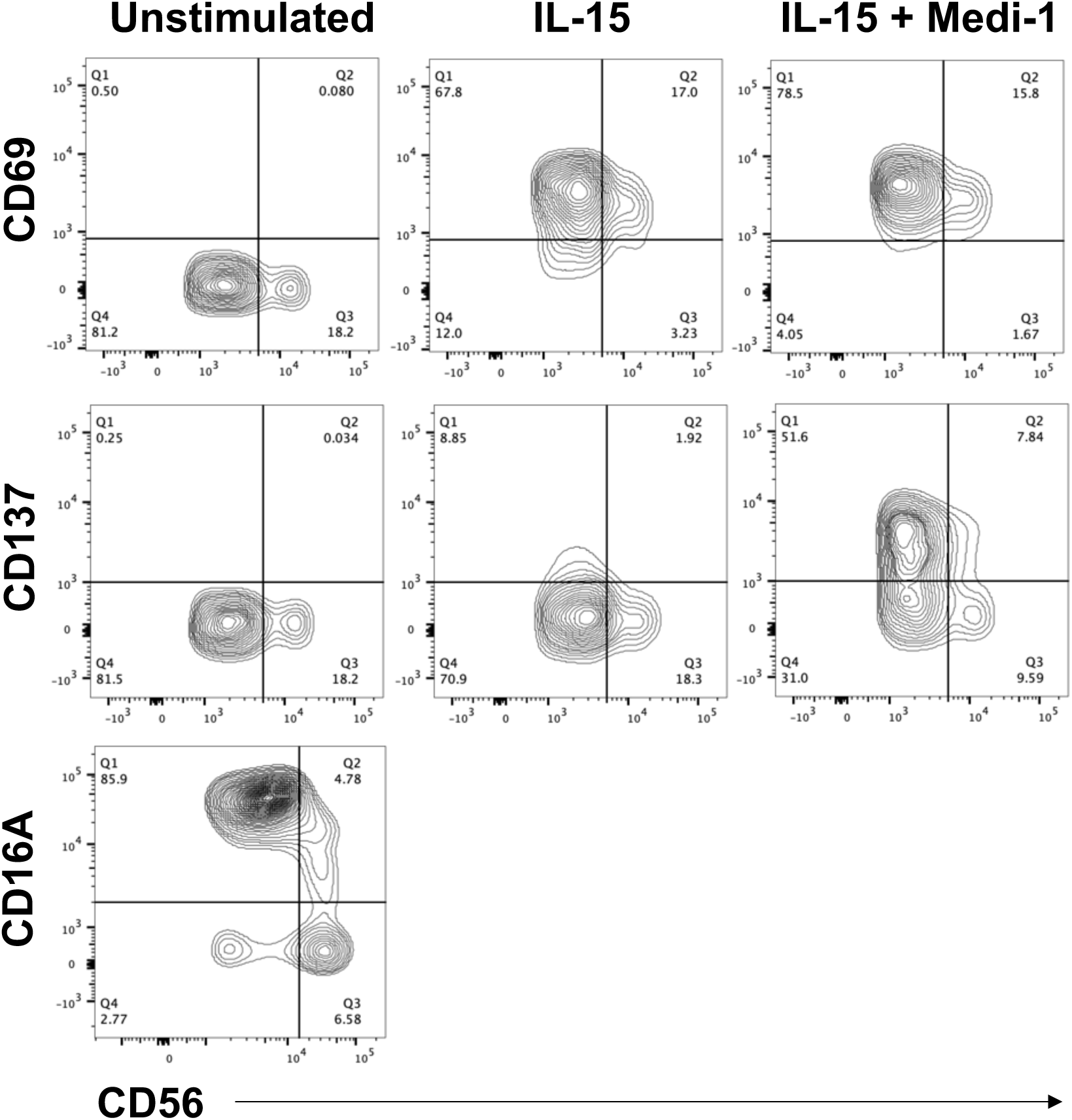
CD137 upregulation primarily occurs by CD56^dim^ versus CD56^bright^ NK cells treated with IL-15 and Medi-1. PBMCs were cultured for 24 hours in the presence or absence of IL-15 (10 ng/ml) +/- Medi-1 (5 µg/ml). CD56^+^ CD3^−^ NK cells were analyzed for their expression levels of the indicated markers by flow cytometry. Data are representative of 3 independent experiments.

**Supplemental Figure 5.**
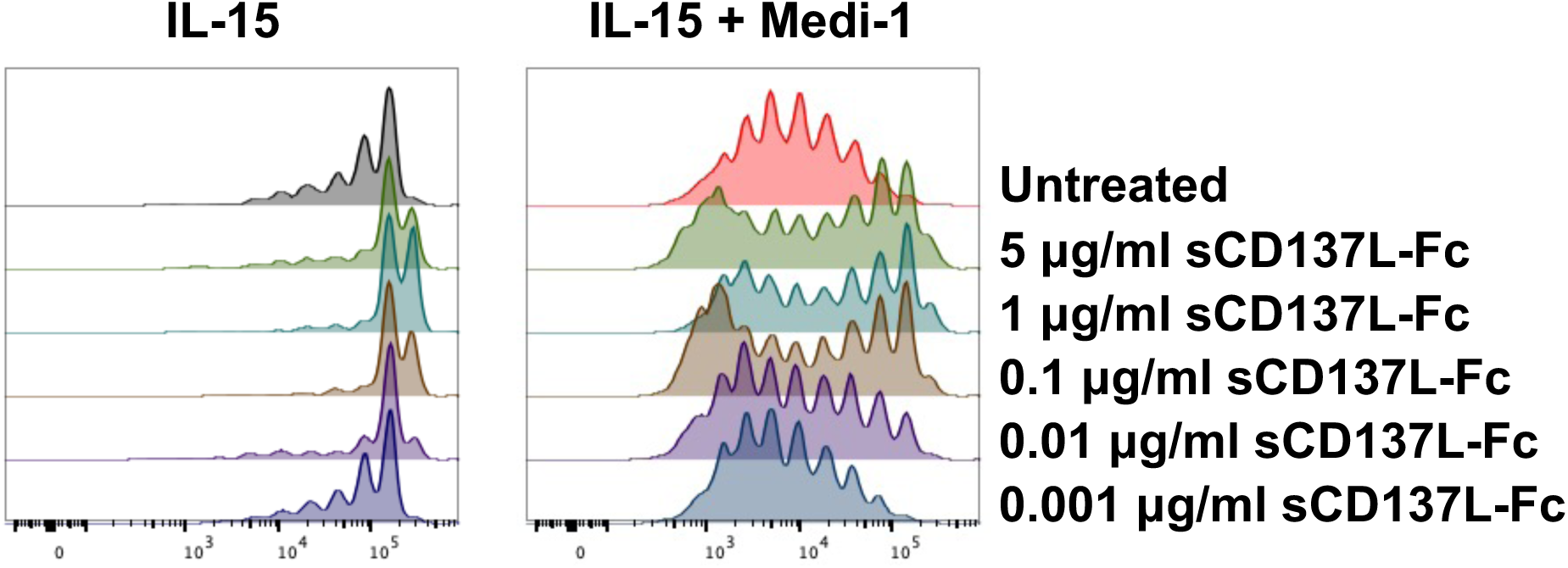
Titration of sCD137L-Fc. PBMCs were labeled with CTV and cultured for 7 days with IL-15 (10ng) +/- Medi-1 (5 µg/ml) +/- sCD137L-Fc fusion protein at the indicated concentrations. CD56^+^ CD3^−^ NK cells were analyzed for CTV dilution by flow cytometry. Data are representative of 3 independent experiments.

